# Restricted transmission of *Xanthomonas oryzae pv. oryzae* from rice roots to shoots detected by a rapid root infection system

**DOI:** 10.1101/2025.08.13.670017

**Authors:** Laura Redzich, Yugander Arra, Eliza P.I. Loo, Wolf B. Frommer

## Abstract

*Xanthomonas oryzae* pv. *oryzae* (*Xoo*), the causal agent of bacterial blight in rice, is primarily studied in the context of foliar infections. However, infected stubble and irrigation water may serve as reservoirs and be responsible for seedling stage root infections in the field, especially during transplanting. Here, we established a coleoptile crown root-based infection protocol to investigate whether gene-for-gene interactions between *Xoo* TAL effectors and *SWEET* sucrose uniporter susceptibility genes occur in the root xylem, and whether the disease can propagate from roots to seedling shoots. Using translational SWEET11a-GUS reporter lines under control of the native SWEET11a promoter, we observed progressive infection as indicated by accumulation of SWEET11a-GUS fusion protein in infected coleoptile crown roots. However, we did not detect progression of GUS accumulation beyond the coleoptile node, nor did we detect blight symptoms on the young leaves. Notably, the xylem, at least during early stages of infection remained functional as shown by Rhodamine B tracer, consistent with transfer of xylem constituents via living cells at the coleoptile node that did not allow bacteria to pass. Furthermore, the root infection protocol is a ∼4x faster compared to standard leaf-clipping assays (roots assay: 11 days from sowing, compared to 39 days for clip infection), enabling more rapid assessment of TAL effector repertoire and plant defense responses with translational SWEET-GUS reporter lines. Our findings expand our understanding of *Xoo* infection routes and provide a valuable tool for resistance testing and pathogen surveillance.

## Introduction

Rice serves as a crucial staple food for developing countries in Asia and Africa (Balasubramanian *et al*., 2007). Bacterial blight, caused by the bacterial pathogen *Xanthomonas oryzae* pv. *oryzae* (*Xoo*), ranks among the most detrimental rice diseases (Liu *et al*., 2014; Savary *et al*., 2019; Swings and Civerolo, 1993). Surveys of bacterial blight outbreaks prevalently report the blight syndrome on leaves of plants at the tillering stage. A severe form of bacterial blight is *kresek* (Javanese: sound of dead leaves) causing complete wilting and lethality of rice at the seedling stage. *Kresek* was associated to *Xoo* after *kresek* symptoms were first reported in Indonesia and predominantly occurred when seedlings were transplanted from nurseries to the field. (Goto, 1964). The route by which *Xoo* infects seedlings, namely via leaves or roots, had remained an open question. Irrigation water and infected rice stubble left in the field after harvest could serve as a reservoir and later as primary inoculum source during the next cropping season (Nyvall, 1999; Ou, 1985; Niño-Liu *et al*., 2006). Consistent with this observation, *Xoo* had been detected in soil and root samples from leaf infected plants (Ritbamrung *et al*., 2025).

The infection mechanism of *Xoo* in leaves relies on a direct gene-for-gene interaction between bacterial transcription activation like effectors (TALes) and *SWEET* promoters. TALe function act as eukaryotic transcription factors to specifically induce transcription of host *SWEETs* by binding to effector binding elements within *SWEET* promoters (Chen *et al*., 2010; Chen *et al*., 2012). Genome editing of the TALe bindings site in the *SWEET* promoters conferred robust resistance to *Xoo* (Eom *et al*., 2019; Oliva *et al*., 2019). TALe-triggered induction of *SWEET* genes is essential for virulence and occurs gene-for-gene specifically. For example, the TALe PthXo1 from the *Xoo* strain PXO99^A^ activates *SWEET11a*, while AvrXa7 of PXO86 induces *SWEET14* (Oliva *et al*., 2019). The ME2 mutant of PXO99^A^ lacking PthXo1 was avirulent and is typically used as a control (Yang and White, 2004).

The standard approach for *Xoo* pathogenicity and resistance testing in rice both in greenhouses and by breeders in the field is leaf clipping assay (Kauffman, 1973), in which leaf tips are cut with that had previously been dipped in *Xoo* suspensions. The length of lesion from the clipping site measured two weeks post-infection serves as an indicator of the degree of susceptibility/resistance to *Xoo*. Here we explored whether rice seedling roots can be infected to evaluate a potential transmission through the xylem from root to shoot, and to develop a fast and reliable protocol for rapid testing of gene-for-gene interaction between TALe and *SWEETs* in rice using reporter lines containing a translational SWEET11a-β-glucuronidase (GUS) fusion under control of the native SWEET11a promoter for root infections (Eom *et al*., 2019). We observed TALe specific induction of *SWEET11a* by the strain PXO99^A^. In contrast to mobility of the small molecule fluorescent dye Rhodamine B, no transmission of the bacteria from coleoptile crown roots to the shoot were observed. The new root assay enables rapid testing of strains within two weeks, approximately four times faster compared to the Kauffman assay (Kauffman, 1973).

## Material and Methods

### Plant material and growth conditions

Rice seeds (Table S1) were dehusked and then sterilized on an orbital shaker at 180 rpm in 15 ml Falcon tubes first for 2 min in 75 % ethanol and then for 5 min in 50 % of a commercial bleach product (2.8 g sodium hypochlorite/ 100g water). Seeds were dried on sterile filter paper and transferred into Magenta™GA-7 boxes containing ½ salt strength MS medium (2.2 g Murashige Skoog Medium (Murashige and Skoog, 1962), 10 g sucrose, 8g Phytagel per liter, pH 5.8) containing 50 mg/L hygromycin B (Table S1) using serological tweezers under aseptic conditions. Seedlings were grown in a 16 h day/8 h night regime with photosynthetic photon flux densities (PPFD) of 200 µmol m^-2^ s^-1^ [PPFD-blue: 40, PPFD-green: 80, PPFD-red: 70 µmol m^-2^ s^-1^] at 27°C, 80 % humidity in CLF PlantClimatics chambers (model: CU41L5).

### *Xanthomonas oryzae* pv. *oryzae* inoculum preparation

*Xoo* cultures (for strains cf. Table S2).) were prepared from glycerol stocks on PSA plates (10 g/L peptone, 1 g/L glutamic acid, 10 g/L sucrose, and 20 g/L agar, pH 7.0). After three days at 28°C in dark conditions, three colonies were picked and placed onto fresh PSA plates. Plates were incubated for additional two days. For the suspension inoculum, *Xoo* was scraped off the plates, resuspended in sterile water and adjusted to OD_600_ 0.5. For the infection, 10 ml of the *Xoo* suspension was transferred to a 50 ml glass beaker.

### Root clip infection

6-day old seedlings were carefully removed from Magenta™GA-7 boxes and rinsed with phosphate-buffered saline (10 mM phosphate, 2.68 mM potassium chloride, 140 m sodium chloride, pH 7.4) to remove residual media. Tips of coleoptile crown roots were cut at 2/3 of their total length (2 cm from the root tip; total length at this stage approximately 3 cm). Tips of cut roots were dipped in *Xoo* inoculum for 10 min (Fig S1) before transfer to DYI hydroponic growth boxes (DYI boxes; empty 3×5 cm 1000 µL pipette tip boxes) filled with Yoshida media (Table S3). Boxes were placed into a PlantClimatics chamber (CU36L) in a 16 h day/8 h night regime at 27°C and 80 % humidity. The evaporation of Yoshida media was monitored daily and the complete media were exchanged accordingly.

### Histochemical GUS assay

For histochemical GUS assays, at 9 am in the morning, seedlings were removed from the DYI boxes, cut approximately 1 cm above the root-shoot interface and collected in 15 ml Falcon tubes containing ice-cold 90 % acetone. Root samples were vacuum infiltrated for 10 min and incubated for 30 min at room temperature. Subsequently, samples were placed in GUS washing buffer followed by staining buffer and were vacuum infiltrated for 10 min each (buffer compositions: Table S4). The enzymatic reaction was stopped after 10 h incubation at 37°C and samples were cleared by replacement of GUS staining buffer with 75 % ethanol for 24 hours.

### Rhodamine B assay

To trace translocation of solutes from roots to shoots, Rhodamine B was used as a small molecule tracer (Wang *et al*., 2022). plants were transferred from the DYI boxes to a double container set-up at 4 dpi (Fig. S2). Both containers were filled with Yoshida media. Either the inner container with infected roots, or the outer container with the newly emerged roots was supplemented with 0.5 mM Rhodamine B (Sigma, 79754). Images were taken 6 days after transfer to the double container set-up (10 dpi).

### Imaging and analysis

Images were taken with a fluorescent Stereo Zoom Microscope (AxioZoom.V16, Zeiss), equipped with a metal halide illuminator (HXP 200C, Zeiss), CMOS cam-era (ORCA-Flash 4.0, Hamamatsu), and PlanNeoFluar objective. For histochemical GUS assays a transparent glass background was used in bright field mode. Rhodamine B was visualized with 63 HE mRFP (ex: 572/25, em: 629/62) on a black background slide. The total length of coleoptile crown roots, the distribution of diX Indigo accumulation, in particular the distance of the diX Indigo front as an indicator of successful infection, and fluorescence were assessed by the segmented line and measurement tools in ImageJ. Analysis was carried out in R (4.05) and RStudio (1.2) employing the R packages tidyverse, ggpubr, ggrepel, lubridate, tigris, sf, dyplr, gapminder, openxlsx, forcats, ggplot2.

## Results

### Successful infection of rice roots by *Xanthomonas oryzae* pv. *oryzae*

To be able to monitor TALe-induced clade III SWEETs, a suite of reporter lines that carry translational SWEET-GUS fusions under control of the respective SWEET promoters had been developed and validated (Eom *et al*., 2019). We recently showed that diX Indigo accumulation in the reporter lines can serve as a specific indicator for TALe activity from different strains. Moreover the reporter lines can be used for monitoring the progression of the *Xoo* infection, in particular at a stage at which *Xoo* had successfully established a Type III Secretion System and injected TALe into host cells (Eom *et al*., 2019; Redzich *et al*., 2025). To explore whether there is a base level of SWEET11a-GUS in different root types, GUS histochemistry was performed on uninfected plants. While crown roots from the coleoptile node with first order lateral roots (hereafter: coleoptile crown roots, Fig S1) did not accumulate diX Indigo, adventitious nodal roots without lateral roots (hereafter: post-embryonic roots) showed SWEET11a-GUS accumulation, indicating a role for SWEET11a in these roots (Fig S3,16). Therefore, only coleoptile roots, but not post-embryonic roots, were used for analysis of potential *SWEET11a* induction in roots infected with PXO99^A^. Coleoptile crown roots were clipped and immersed with the apical region in *Xoo* inoculum for 10 minutes. In contrast to mock of ME2 control strain infections (Yang and White, 2004; Eom *et al*., 2019), PXO99^A^ caused diX Indigo accumulation in the root vasculature (Fig 1A, B). When roots of SWEET11a-GUS expressing lines were infected with *Xoo* strain PXO86, which carries AvrXa7 targeting the promoter of *SWEET14*, but which has no TALe to induce *SWEET11a*, no diX Indigo accumulation was detectable, demonstrating that PXO99^A^ must have been able to successfully been able to inject pthXo1 into host cells and to induce SWEET11a. Notably, the front of *SWEET11a* induction, represented by diX Indigo, advanced ∼0.4 cm per day after infection (Fig 1C), indicating that the bacteria progressively infected the root at this velocity. The distance of the diX Indigo front as an indicator of the progression of the infection from the clip site progressed further between 2 and 4 dpi (days post infection), while no further progression was observed at the shoot-root axis (Fig S4). In no case did we diX Indigo in the shoot (except for wound-induced staining at the cut site). Thus, in comparison to the leaf clipping method performed on 4-week-old plants and lesion scoring two weeks dpi, the root infection protocol is ∼4x faster (11 days from germination to evaluation: 6 days seedling growth; 4 days infection; 1 day for GUS histochemistry).

**Figure 1.**
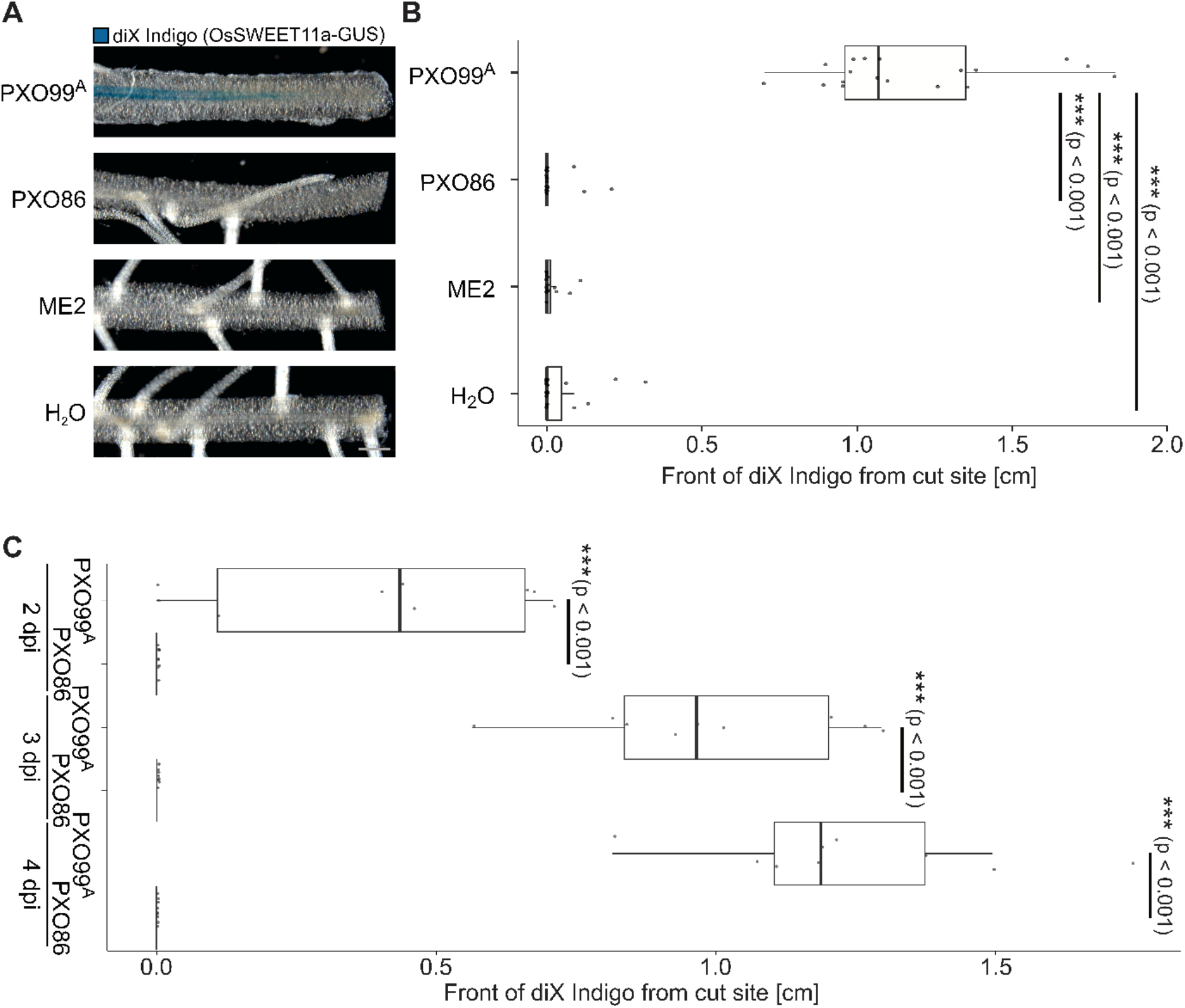
*Xoo* strain-specific induction of GUS activity in SWEET11a-GUS reporter lines after infection of coleptile crown roots. diX Indigo accumulated after infection with PXO99^A^, but not after mock, ME2 or PXO86 infections. **A)** diX-Indigo accumulation at (4 dpi) after infection with PXO99^A^, PXO86, ME2 or mock (water) in the root vasculature observed with the Zeiss Axiozoom V16 and processed in ImageJ (bar: 0.5 mm). Representative images from 50 roots per treatment from three independent experiments performed with two independent pSWEET11a:SWEET11a-GUSplus lines (independent transformants: line #8 and #10)(Eom *et al*., 2019)(Table S2.1). **B)** Progression of the infection as indicated by the distance of the diX Indigo front from the cutting site in coleoptile crown roots treated with PXO99^A^, PXO86, ME2, or H_2_O at 4 dpi. Measurements were performed in ImageJ with the Segmented Line Tool. Data processing performed in R. **C)** Time-dependent progression of the infection as indicated by the distance of the diX Indigo front from the cutting site in coleoptile crown roots 2, 3 and 4 dpi after PXO99A infection. PXO86 served as a control. Repeated independently three times with comparable results.

### Restricted vascular progression of *Xoo* to aerial tissue in rice

*Kresek* symptoms prevalently cause wilting of seedling coleoptile and leaves. One may speculate that wounds at roots, e.g. during transplanting, could serve as an entry route for infection. Plant debris, irrigation water and soil may serve as reservoirs for *Xoo*. Consistent with the inability of *Xoo* to progress beyond the coleoptile node, we did not observe disease/ *kresek* symptoms in the seedlings using the same infection method (Fig S5). The inability to enter the shoot might be due to rapid blockage of the xylem in response to wounding. To evaluate whether the xylem is blocked, a double-container set up was generated in which infected coleoptile crown and non-infected post-embryonic roots were immersed in separate compartments with hydroponic media. Rhodamine B was used as fluorescent marker for xylem mobility (Wang *et al*., 2022). Rhodamine B was detected x days after addition Roots clipped and transferred into the double container set-up 4 dpi (Fig 2). Ten days post infection, the progression of Rhodamine B was evaluated in leaves 5 cm above the coleoptile node with fluorescence microscopy. Post-embryonic roots showed translocation of Rhodamine B into major and minor veins of the leaf, demonstrating functional xylem translocation. The inability of bacteria to pass the coleoptile node must thus be due toother reasons. Bacterial infections were apparently able to block the xylem transport 4-10 days after infection, Rhodamine B translocation was undetectable, indicating that at later stages of infection, xylem transport is blocked.

**Figure 2.**
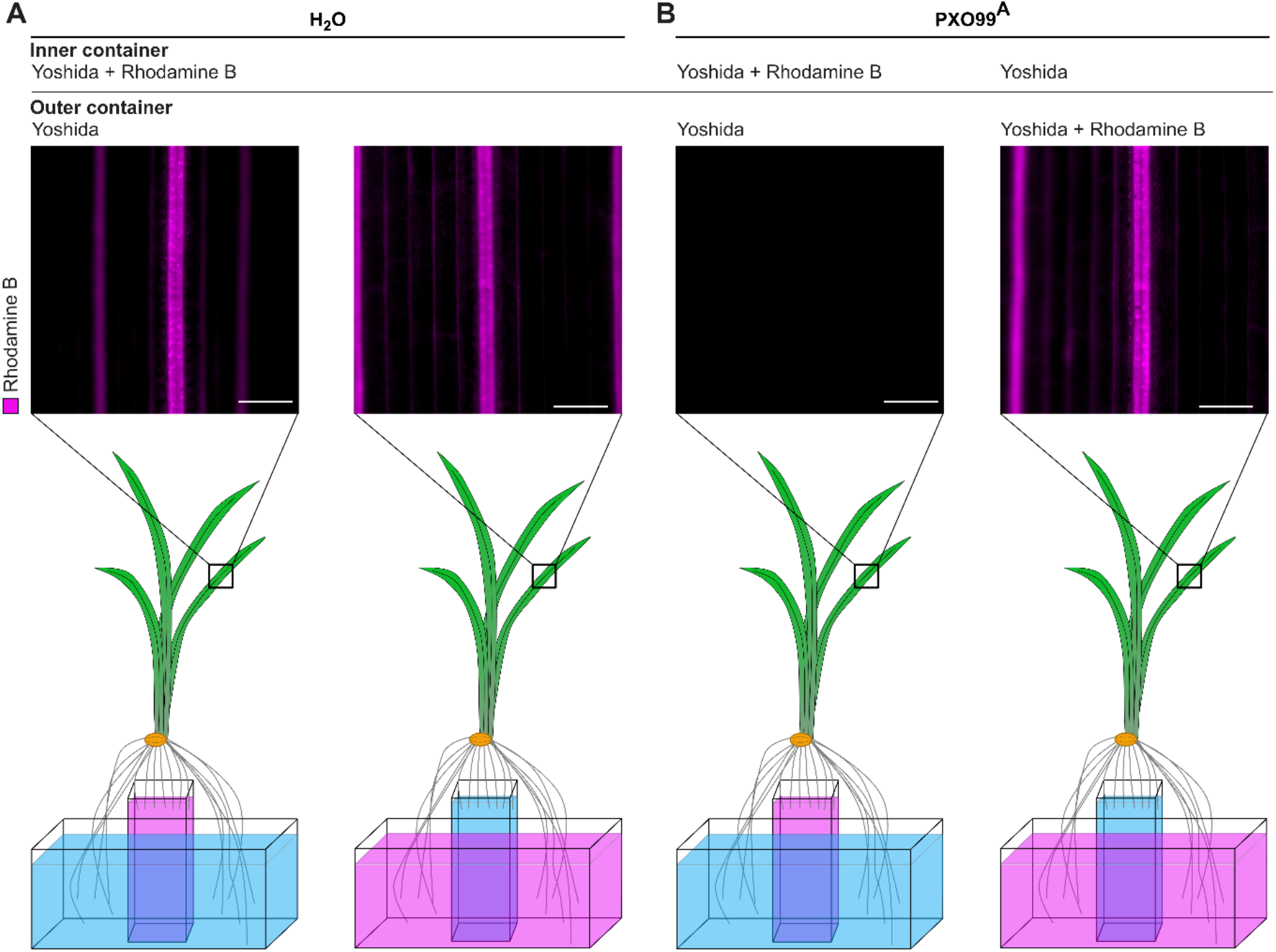
Root-to-shoot xylem connectivity in seedlings before and after *Xoo* infection assayed using Rhodamine B as a tracer. **A)** Rhodamine translocation from coleoptile crown roots (left) or postembryonic roots (right) in uninfected seedlings. 10-day-old seedlings were transferred to a double container set up 4 days after clipping coleoptile crown roots. Roots were immersed in Yoshida medium (blue) or Yoshida medium supplemented with Rhodamine B (purple), respectively. Bottom shows cartoon of setup, top shows Rhodamine B fluorescence in the veins of seedling leaves 5 cm above the coleoptile node acquired using a Zeiss Axiozoom V16 and processed in ImageJ and Omero (n=3). **B)** Rhodamine translocation from coleoptile crown roots (left) or postembryonic roots (right) in *Xoo*-infected seedlings. 10-day-old seedlings were transferred to a double container set up 4 days after clipping coleoptile crown roots and infection by *Xoo* for 10 minutes. Scale bar: 1 mm. Repeated independently three times with comparable results.

## Discussion

*Xoo* infects the xylem and its virulence depends critically on the induction of host SWEET sucrose transporters in the xylem parenchyma by bacterial TALe (Eom *et al*., 2019; Oliva *et al*., 2019; Chen *et al*., 2010; Chen *et al*., 2012; Schepler-Luu *et al*., 2023). *Xoo* acquires host-derived sucrose in the xylem with the help of transporters and enzymes of the *Sux* gene cluster (Zöllner *et al*., 2025). Importantly, key virulence functions and virulence depend on the ability to utilize sucrose (Zöllner *et al*., 2025). At present, six different TALe have been identified that are utilized by different *Xoo* strains and which each target a specific SWEET paralog in rice (Oliva *et al*., 2019). The targeted SWEETs are part of a specific clade of six sucrose uniporters(Wu *et al*., 2022). While all six clade 3 SWEETs can function as susceptibility genes for *Xoo*, only three have been found so far to be targeted by *Xoo* strains (Streubel *et al*., 2013). Breeding of resistance and genome editing are effective tools for controlling bacterial blight, however new strains have either emerged that break resistance or disease outbreaks were caused by inadvertent introduction of strains from other areas or even continents (Raveloson *et al*., 2023; Schepler-Luu *et al*., 2023; Sciallano *et al*., 2023). It is thus critical to develop tools that enable rapid identification of SWEET genes targeted by emerging *Xoo* strains as a basis for rapid development of new lines that carry suitable promoter edits on SWEET promoters (Eom *et al*., 2019; Oliva *et al*., 2019; Schepler-Luu *et al*., 2023).

To enable rapid identification of which SWEET is targeted by an emerging *Xoo* strain, we developed a SWEET^R^ kit, which included full gene translational SWEET-GUS reporter lines (Eom *et al*., 2019; Wu *et al*., 2022). These reporter lines also enable the monitoring of the infection process using the Kauffman leaf clipping assay, which is typically performed on four- to six-week-old plants and recording symptoms 14 days post infection (Redzich *et al*., 2025). Because plants are most vulnerable towards *Xoo* within 21 days after germination, it is necessary for reliable infection scoring, that plants are infected at the maximum tillering stage at about four weeks after germination (Kauffman, 1973). If plants are infected at earlier developmental stages using the leaf clipping method, *kresek* symptoms can bias the lesion scoring results towards higher susceptibility (Mew *et al*., 1979).

The aim of this study was fourfold: (i) evaluate whether *Xoo* can infect wounded roots, (ii) explore whether the infection of roots depends, similar as for leaves, on the induction of SWEETs, (iii) whether the bacteria can migrate from roots to shoots, and (iv) develop a faster test system to identify which SWEET is targeted by an emerging *Xoo* strain.

i. Using two rice lines carrying translational SWEET11a-GUS fusion driven by the *SWEET11a* promoter, we found that *Xoo* can infect the root xylem in the same manner as the leaf xylem after wounding (root tip clipping). The reporter lines also enabled monitoring the progression of the infection in roots over time. The assay could only be used in coleoptile crown roots, since postembryonic roots had base levels of SWEET11a and thus induction could not be observed against the background.
ii. The induction of SWEET11a was specific for the strain PXO99^A^, which produces the TALe PthXo1 that targeted the SWEET11a promoter.
iii. Interestingly, SWEET induction occurred only in the roots and did not progress beyond the coleoptile node. We also did not observe any symptoms in the shoot post root infection, indicating that *Xoo* cannot pass from coleoptile crown roots to the shoot xylem. In contrast, the small molecule dye Rhodamine B was effectively translocated to the shoot. Nodal translocation zones contain three distinct vascular bundles: enlarged vascular bundles (EVBs), transit vascular bundles (TVBs) and diffuse VB (DVBs, (Yamaji and Ma, 2014). *Xoo* would need to either translocate via the nodal vascular anastomosis to the DVB, or exit xylem vessels of EVBs to living xylem transfer cells to the DVB connecting to the upper node. Alternatively, *Xoo* would need to travel along the TVB to the multitude of xylem vessels of EVBs to reach the leaf (Yamaji and Ma, 2014; Schwab *et al*., 2016). Likely due to the discontinuity of the vessel and the transfer of xylem constituents via living cells, the dye could be transferred, but not the *Xoo* cells. Because *Xoo* did not translocate to leaves, likely the infection was restricted to roots and systemic infection as described for *kresek* did not occur. The results shown here indicate that *kresek* symptoms are not caused by plant entry of *Xoo* by root wounding after transplantation, but rather are transmitted via shoots. We also did not observe *kresek* symptoms when all roots had been clipped. However, that the exposure to *Xoo* in our experiments was limited to 10 minutes, it is conceivable that long term exposure may also lead to transmission via newly emerging roots. Of note, at later stages of infection, translocation of Rhodamine B ceased, indicating that the bacteria blocked xylem flow either due to clogging by biofilm, tylosis or due to xylem cell death.
iv. Leveraging the suite of SWEET-GUS reporters from presented in Eom *et al*. 2019, the root infection assay of translational SWEET-GUS reporter lines allows for four-times faster screening for *SWEET* induction, i.e. results can be obtained within 11 days. Full development of the assay for all six SWEETs will require testing of the base levels of the other five clade 3 SWEETs in the different root types.

The root infection protocol developed here provides a fast method to screen naturally occurring TALe variants and support pathogen surveillance. We surmise that after analysis of the base levels of SWEET-GUS reporter activity of all six clade 3 SWEET -GUS reporter lines the test could be expanded to rapidly evaluate which SWEET is targeted by a novel TALe of an emerging strain (Eom *et al*., 2019; Wu *et al*., 2022). Furthermore, the infection protocol might be a promising tool to test TALe functionality and plant defense responses.

## Data availability

Raw data are available at https://doi.org/10.60534/7s2cg-r2j06

## Contributions

LR, YA, EL and WF conceived of the study. LR performed clipping infection, histology assays and Rhodamine B experiments, analysis and wrote the manuscript. YA established rice root infection protocol, performed root infections to trace *Kresek* phase. EL supervised team. LR, YA and WF analyzed the data and wrote the manuscript.

## Acknowledgments

This work was supported by grants from Deutsche Forschungsgemeinschaft to WBF (DFG, German Research Foundation) - Collaborative Research Center SFB1535, project ID 458090666/CRC1535/1 the Alexander von Humboldt Professorship (WBF); funds from Germany’s Excellence Strategy – EXC-2048/1 – project ID 390686111 (Deutsche Forschungsgemeinschaft, CEPLAS) to WBF and LR; fellowships to LR from Heinrich-Böll-Stiftung, and to YA by the Alexander von Humboldt Foundation, and support from the Director, CSIR-Central Institute of Medicinal and Aromatic Plants, India.

