## Supplementary material for "Restricted transmission of *Xanthomonas oryzae pv. oryzae* from rice roots to shoots detected by a rapid root infection system": Fig S1

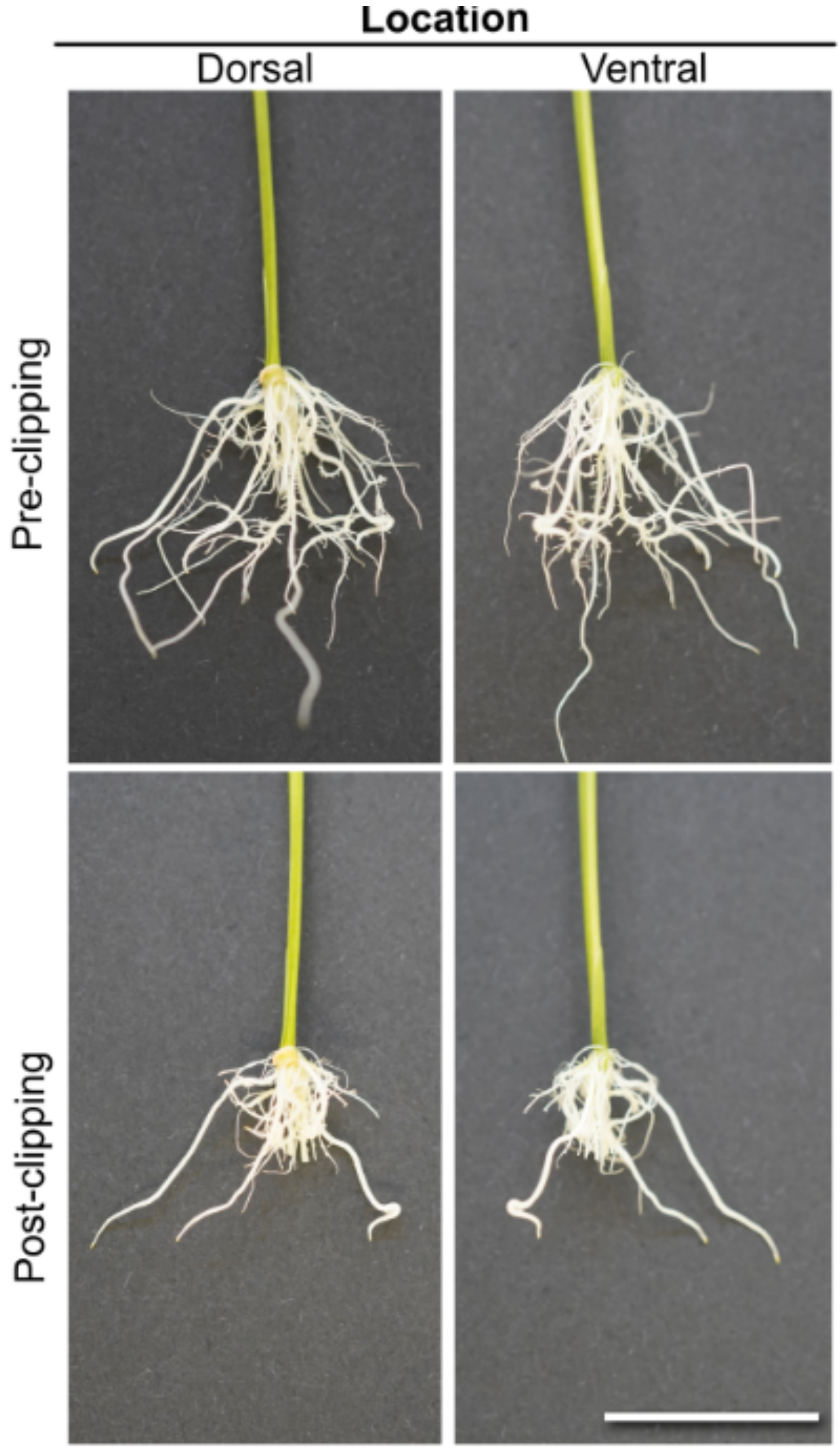


**Supplemental Figure S1. Clipping of coleoptile crown roots in root infection protocol.** Coleoptile crown roots with first order lateral roots emerge from coleoptile node. Post-embryonic adventitious crown roots without lateral roots emerging from the first node were not clipped. Tips of coleoptile crown roots were cut at 2/3 of their total length (2 cm from the root tip; total length at this stage approximately 3 cm. Root systems shown before and after clipping. Nomenclature as proposed by the International Society of Root Research (Freschet *et al.*, 2021).Ventral (left) and dorsal (right). Scale bar: 1 cm.

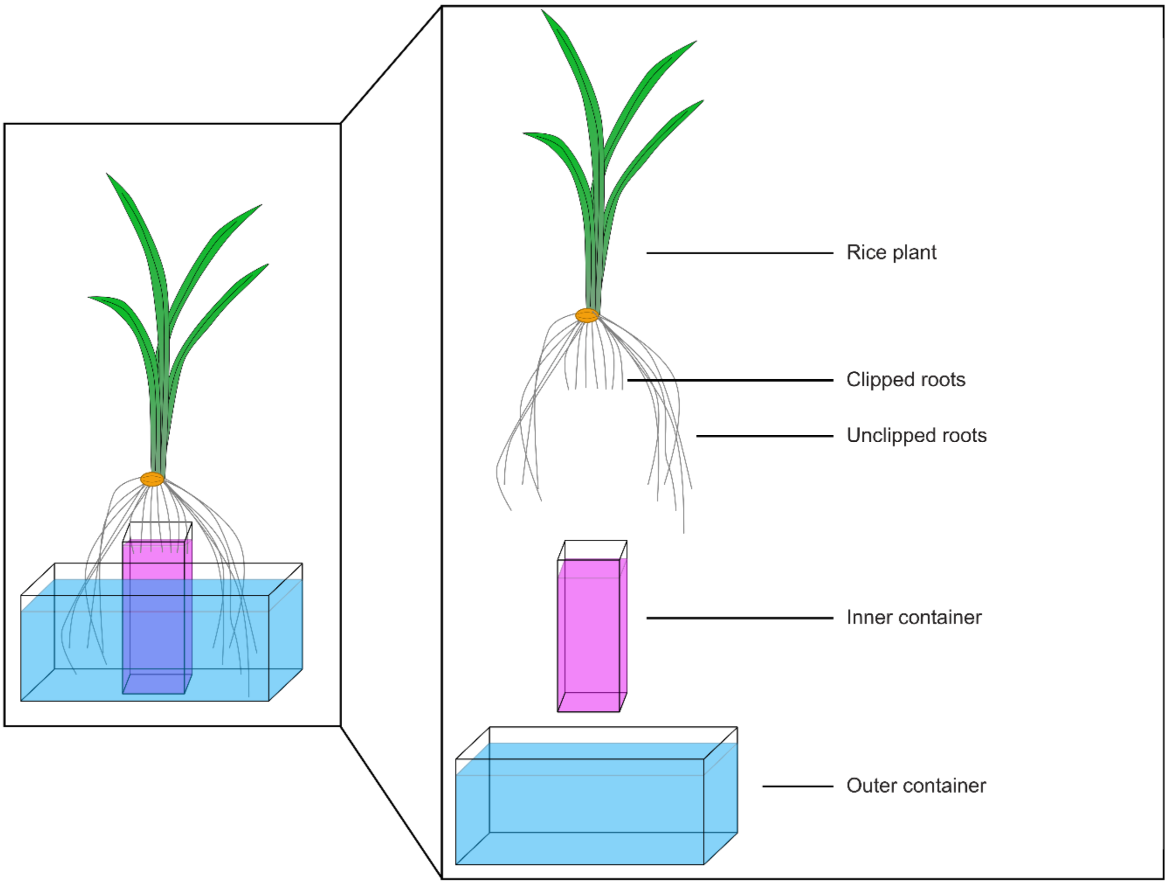


**Supplemental Figure S2.** Double container set-up to track translocation of solutes clipped and non-clipped roots to leaf vasculature. A 50 mL plastic beaker (inner container) was glued onto the bottom of a 3x5 cm 1000 µL pipette tip box (outer container). By the addition of Rhodamine B to the outer or inner container, translocation via unclipped or clipped roots were investigated. Plants were stabilised with tape and wooden toothpicks.

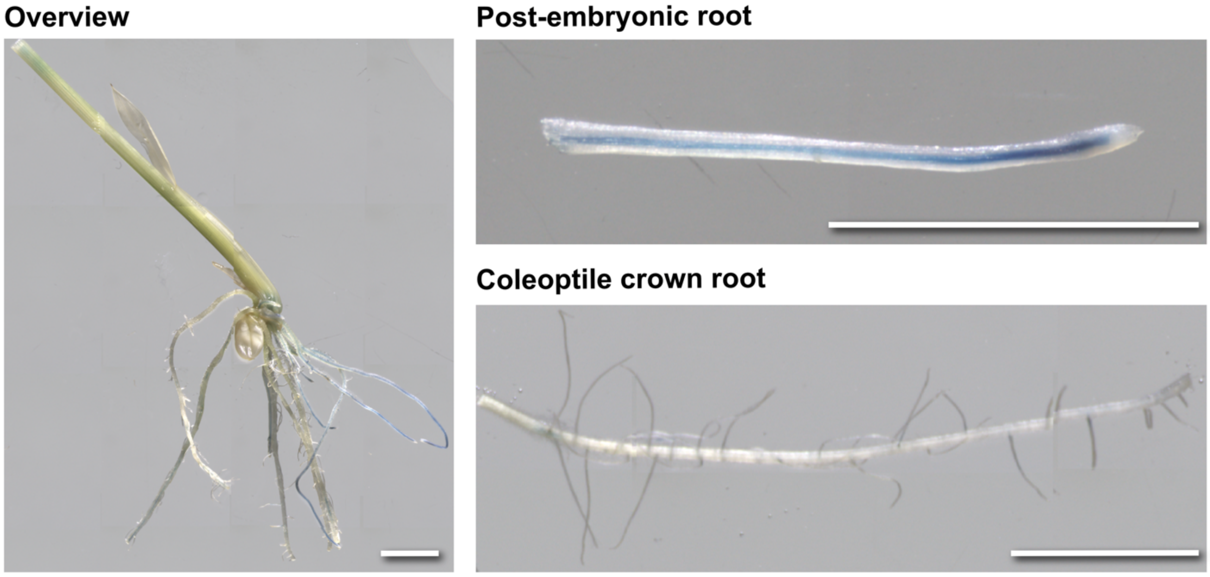


**Supplemental Figure S3. Accumulation of diX Indigo in uninfected root system.** Coleoptile crown roots with first order lateral roots did not accumulate diX Indigo. Post-embryonic adventitious crown roots without lateral roots accumulated diX Indigo independent of TALe induction. Scale bar: 0.5 cm.

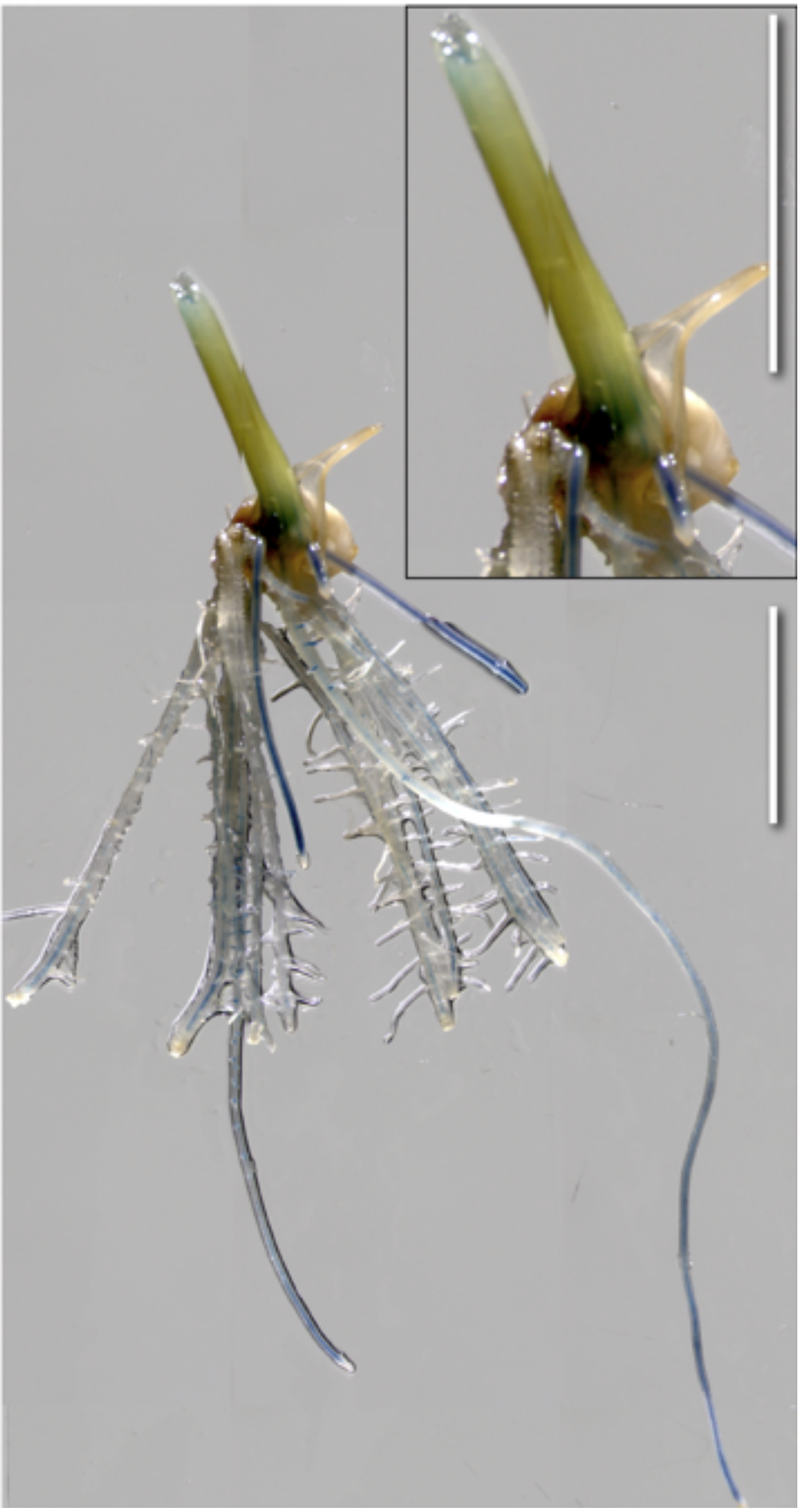

**Supplemental Figure S4. diX Indigo accumulation is restricted to roots.** Coleoptile crown roots with first order lateral roots and post-embryonic crown roots accumulated diX Indigo. diX Indigo was not observed in the shoot (except at the cutting site). Scale bar: 1 cm.

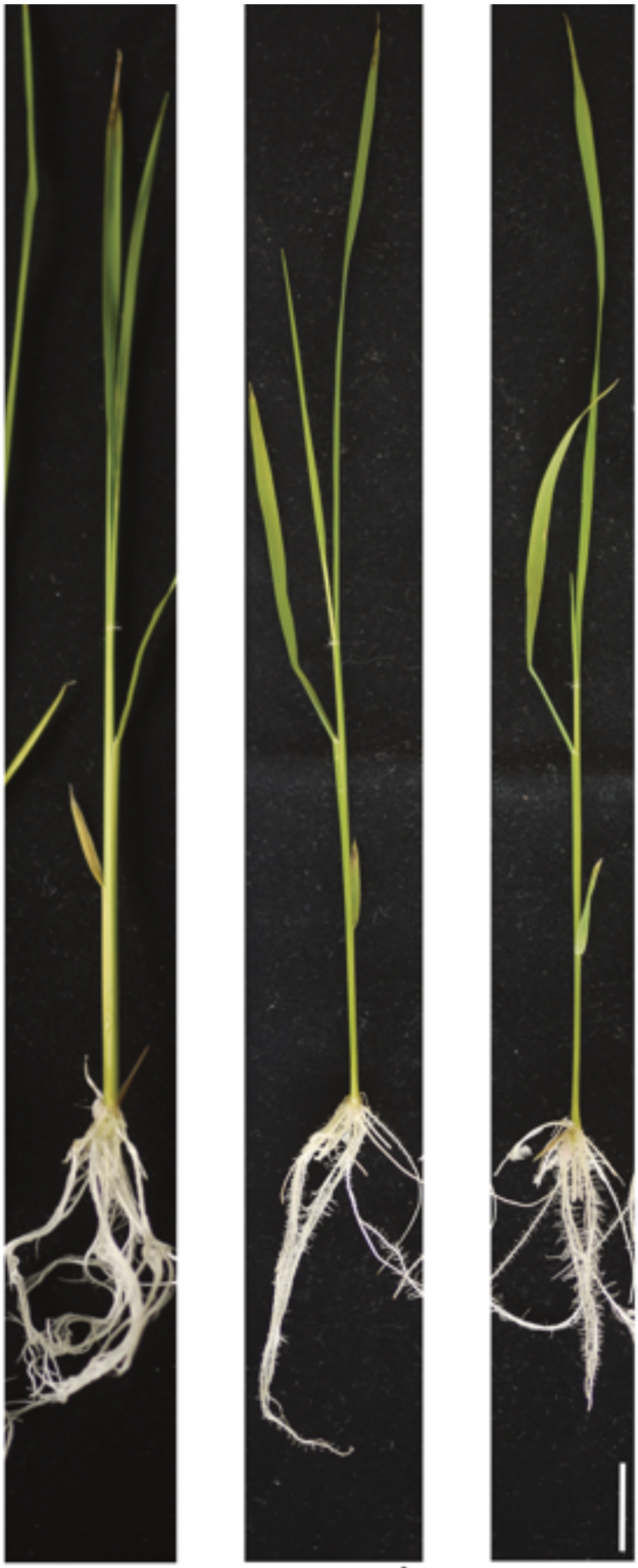


**Supplemental Figure S5. Phenotype of seedlings seven days post infection.** Roots were clip infected with PXO99$^A$, PXO86 and mock treated with $H_2O$. *Kresek* symptoms were not observed. Scale bar: 2 cm.

**Supplemental Table S1. Overview of rice lines.** All lines were retrieved from the rice seed stocks and tested for Hygromycin B resistance.

| Name | Transformation Event | Characteristics | Generation | Resistance Ratio |
|---|---|---|---|---|
| B01069 | [8-3] | pSWEET11a:SWEET11a-GUSplus | T4 | 1.0 |
| B01075 | [10-2] | pSWEET11a:gSWEET11a-GUSplus | T4 | 1.1 |

**Supplemental Table S2. Overview of *Xoo* strains.** All strains were retrieved from the -80 °C glycerol stocks.

| Strain | TALE | Targeted EBE sequence | Targeted *SWEET* |
|---|---|---|---|
| PXO86 | AvrXa7 | ATAAACCCCCTCCAACCAGGTGCTAA | *SWEET14* |
| PXO99[A] | PthXo1 | GCATCTCCCCCTACTGTACACCAC | *SWEET11a* |
| ME2 | - | | non-virulent |

**Supplemental Table S3. Stock solutions for Yoshida media**. For 4 liters of working solution, add 2.5 ml of each stock solution and adjust to pH 5.8 with KOH.

| Stock order | Element | Chemical | Preparation (g/10 L $H_2O$) | Remarks |
|---|---|---|---|---|
| 1 | N | $NH_4NO_3$ | 914 | Final volume of 10 L |
| 2 | P | $NaH_2PO_4.H_2O$ | 403 | Final volume of 10 L |
| 3 | K | $K_2SO_4$ | 717 | Final volume of 10 L |
| 4 | Ca | $CaCl_2$ | 886 | Final volume of 10 L |
| 5 | Mg | $MgSO_4.7H_2O$ | 3240 | Final volume of 10 L |
| **Micronutrients** | | | | |
| 7 | Mn | $MnCl_2.4H_2O$ | 15 | Prepare separately, mix and adjust final volume of 10 L |
| 8 | Mo | $(NH_4)_6.Mo_7O_{24}.4H_2O$ | 0.74 | |
| 9 | B | $H_3BO_3$ | 9.34 | |
| 10 | Zn | $ZnSO_4.7H_2O$ | 0.35 | |
| 11 | Cu | $CuSO_4.5H_2O$ | 0.31 | |
| 12 | Fe | Fe-Na-EDTA | 104 | |

**Supplemental Table S4. Recipes for solutions for GUS histochemistry**

**Washing buffer**

| Component | Final concentration | For 50 ml |
|---|---|---|
| 0.5 M EDTA | 10 mM | 1 ml |
| 100 mM phosphate buffer, pH 7 | 50 mM | 25 ml |
| 10 % triton X-100 | 0.1 % | 0.5 ml |
| 50 mM potassium ferrocynide | 1 mM | 1 ml |
| 50 mM potassium ferricynide | 1 mM | 1 ml |
| Methanol | 20 % | 10 ml |
| Sterile water | | Ad 50 ml |

**X-Gluc staining buffer**

| Component | Final concentration | For 50 ml |
|---|---|---|
| 0.5 M EDTA | 10 mM | 1 ml |
| 100 mM phosphate buffer, pH 7 | 50 mM | 25 ml |
| 10 % Triton X-100 | 0.1 % | 0.5 ml |
| 50 mM potassium ferrocyanide | 1 mM | 1 ml |
| 50 mM potassium ferricynide | 1 mM | 1 ml |
| Methanol | 20 % | 10 ml |
| 100 mM X-Gluc | 2 mM | 1 ml |
| Sterile water | | Add 50 ml |

**100 mM Phosphase buffer, pH 7**

| Component | Volume |
|---|---|
| 0.5 M sodium phosphate dibasic ($Na_2HPO_4$) | 3 ml |
| 1 M sodium phosphate monobasic ($NaH_2PO_4$) | 1 ml |

**100 mM X-Gluc solution**

| Component | Final concentration | For 50 ml staining solution |
|---|---|---|
| X-Gluc | 100 mM | 50 mg |
| DMSO | | 1 ml |
